# Toxicity drives facilitation between four bacterial species

**DOI:** 10.1101/605287

**Authors:** Philippe Piccardi, Björn Vessman, Sara Mitri

## Abstract

Competition between microbes is extremely common, with many investing in a wide range of mechanisms to harm other strains and species. Yet positive interactions between species have also been documented. What makes species help or harm each other is currently unclear. Here, we studied the interactions between four bacterial species capable of degrading Metal-Working Fluids (MWF), an industrial coolant and lubricant, which contains growth substrates as well as toxic biocides. We were surprised to find only positive or neutral interactions between the four species. Using mathematical modeling and further experiments, we show that positive interactions in this community are likely due to the toxicity of MWF, whereby each species’ detoxification benefited the others by facilitating their survival, such that they could grow and degrade MWF better when together. The addition of nutrients, the reduction of toxicity or the addition of more species instead resulted in competitive behavior. Our work provides support to the stress gradient hypothesis by showing how harsh, toxic environments can strongly favor facilitation between microbial species and mask underlying competitive interactions.

## Introduction

A microbial cell living in the human gut, in the soil or in a biofuel cell is typically surrounded by cells of its own as well as other strains and species. The way in which it interacts with other community members is key to its growth and survival, and ultimately, to the stability and functioning of the community as a whole (1, 2). Being able to predict community dynamics and functioning over ecological and evolutionary time-scales is not only fundamentally interesting, but can also help develop therapies for microbiome dysbiosis or augment soil to improve agricultural productivity (2–7).

A central question in studying microbial interactions is whether community members cooperate or compete with one another (8–10). Stable cooperation that evolves in two interacting species because of their benefit to one another (9) is only expected under highly restrictive conditions (11, 12), with few documented examples (13). Facilitation (14) is more prevalent since it encompasses cooperation as well as commensalism, where one species accidentally benefits from another, for example by cross-feeding off its waste products (15–19). It appears, however, that microbial life is mostly competitive: Microbes have evolved a great number of ways to harm other strains and species (20). For example, 25% of gram-negative bacteria possess genes coding for a Type VI Secretion System (21), while 5–10% of actinomycete genomes code for secondary metabolites (22). Such aggressive behavior likely evolved due to competition for available resources, be they nutrients, oxygen or space. Our base expectation is therefore that microbial species will tend to compete (9, 11).

However, whether species help or harm each other appears to depend on environmental gradients (23–29). The Stress Gradient Hypothesis (SGH, (30)), predicts that positive interactions should be more prevalent in stressful environments, while permissive environments should favor competition. The hypothesis has only rarely been tested in microbial communities (23, 29, 31, 32) and the studies that have tested it involve either species whose interactions have been genetically engineered (29), theoretical work (32), or communities containing many species (23, 31), where it is difficult to quantify individual species abundances and their interactions, and to understand why observations are in line with the SGH.

To fill this gap, here we used a synthetic community composed of four bacterial species that has been applied to the bioremediation of highly alkaline and polluting liquids used in the manufacturing industry called Metal-Working Fluids (MWF) (33–35). MWFs contain chemical compounds that are rich nutrient sources for bacteria, such as mineral oils and fatty acids (36), as well as biocides that inhibit microbial activity (35, 37). The four species – *Agrobacterium tumefaciens*, *Comamonas testosteroni*, *Microbacterium saperdae*, and *Ochrobactrum anthropi* – were previously isolated from waste MWF and selected based on their ability to individually survive or grow in MWF (34). The synthetic community was shown to degrade the polluting compounds in MWF more efficiently and reliably than a random community (34, 38). This community in its defined chemical environment, represents a tractable model system for exploring how abiotic and biotic interactions shape the ecological dynamics of microbial communities. By quantifying MWF degradation efficiency and mapping it to species composition and their interactions, this model system can also help answer another key question in microbial ecology: how do inter-species interactions affect ecosystem functioning?

Below, we show that when growing in MWF, facilitation dominates interactions between these four species, and that this is likely due to the toxicity of MWF. By making the environment more permissive, we further show that interactions become competitive, in a pattern that is consistent with the SGH. In turn, degradation efficiency only improves with community size when the environment is toxic and interactions are positive. Our experiments shed light on how nutrient and toxicity gradients modulate interactions between species and community functioning.

## Results

### Facilitation dominates the community in MWF

We first characterized the effect of each species in the MWF community on the others. The four species were incubated alone (mono-culture) or in combination with a second species (pairwise co-culture) in shaken flasks containing MWF medium over 12 days (see Methods). The inoculum volume for each species was held constant across all conditions, i.e. the total was higher in co-cultures. In mono-culture, *C. testosteroni* was able to survive and grow in MWF, while *A. tumefaciens* survived in some replicates, and *M. saperdae* and *O. anthropi* did not (Fig. 1A-D). Qualitatively similar results were obtained in an independent repeat of the experiment (Fig. S1). We quantified species interactions by comparing the area under the growth curve (AUC) of mono- and pairwise co-cultures and define an interaction as negative or positive if the AUC of the co-culture is significantly smaller or greater than the AUC of the mono-culture, or neutral otherwise (see Methods). Defining interactions by the AUC means that they may vary with the length of the experiment and the inoculum volume, but the measure nevertheless combines growth rate, death rate and final yield in one value. Using this measure, positive interactions dominated the MWF ecosystem (Fig. 1E, 3A, S1). *C. testosteroni* promoted the survival and growth of all other species, while also benefiting significantly from the presence of *A. tumefaciens* and *M. saperdae*. *M. saperdae* and *O. anthropi* also slightly reduced each other’s death rates (Fig. 1C, D). Finally, *A. tumefaciens* rescued *M. saperdae* from extinction (Fig. 1C), but the AUC was not significantly different from *M. saperdae* in mono-culture.

**Fig. 1.**
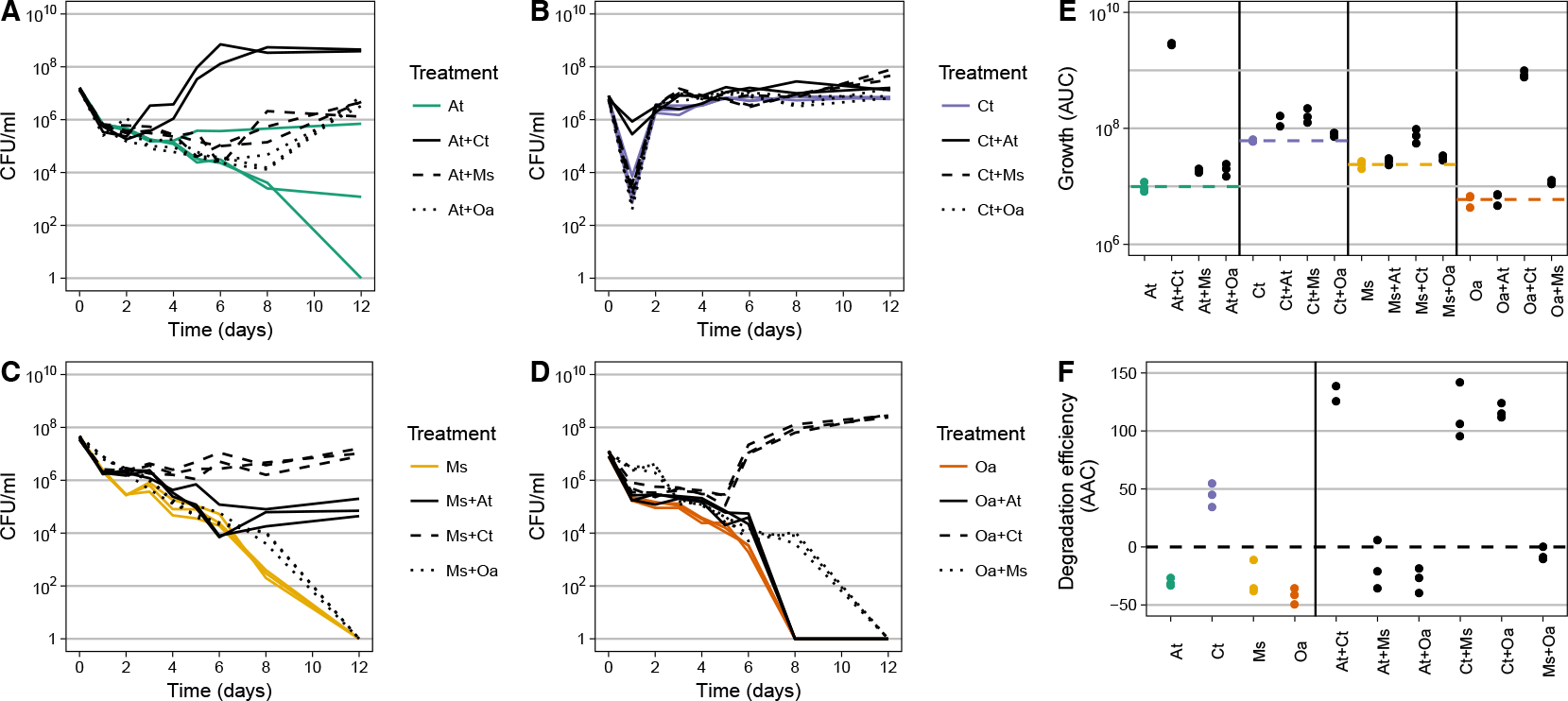
Comparison of mono- and pairwise co-cultures. (**A-D**) Population size quantified in CFU/ml over time for mono-cultures (in color) and pairwise co-cultures (in black) of *A. tumefaciens* (At), *C. testosteroni* (Ct), *M. saperdae* (Ms) and *O. anthropi* (Oa) in panels (**A**) to (**D**), respectively. (**E**) Area under each of the curves (AUC) in panels (**A-D**). Dashed lines indicate the mean of the mono-cultures, shown in color. Statistical significance is calculated based on combined data from this and the repetition experiment (Fig. S1), and shown in Fig. 3 and Table S2. (**F**) Area above the curves (AAC) describing the decrease in Chemical Oxygen Demand (COD, see Methods) over time (i.e. degradation efficiency, Fig. S6A, B). Negative AAC values arise because dead cells increase the COD (Fig. S7). AUC (**E**) and AAC (**F**) correlate positively (Fig. S4).

We wondered whether these positive interactions between species were specific to these four species, which may have adapted to each other’s presence in the past (38). To test for this we grew new isolates that had never previously interacted with our four species, together with *C. testosteroni* and found similar two-way positive effects in MWF (Fig. S2, S3). This suggests that these positive interactions are likely to be accidental rather than having evolved because of their positive effect (facilitation rather than evolved cooperation).

Degradation efficiency in all co-cultures that included *C. testosteroni* showed a higher compared to any of the mono-cultures (Fig. 1F). More generally, degradation efficiency correlated positively with population size (Fig. S4, Spearman’s *ρ* = 0.77, *P* < 10^*−*15^).

### Facilitation is not due to inoculum doubling

In our experimental setup, the total initial inoculated population was larger in co-cultures compared to the mono-cultures. If all four species detoxify and degrade the same exact compounds in MWF, positive interactions could be explained by this larger initial cell density. Alternatively, if species differ in their contribution to detoxification, positive interactions should be maintained even if we keep the initial cell density constant across treatments.

To differentiate between these possible explanations, we repeated the experiment with a constant total inoculum volume across pairwise co-cultures and mono-cultures. All species still grew significantly better in the presence of *C. testosteroni*, and *C. testosteroni* benefited from all others (Fig. S5). However, *M. saperdae* and *O. anthropi* died faster in pairwise co-cultures compared to mono-cultures if their partner was also dying. Worse growth was presumably due to halving the focal species’ inoculum, rather than a real negative interaction between these species pairs. Indeed, doubling the number of cells in mono-culture showed a significant improvement for all species (F-test, df=3, all *P* < 0.015). In other words, even though the starting population size of mono-cultures influences survival, the four species appear to functionally complement each other in facilitating growth and survival in MWF.

Together, these first results appear to contradict the expectation that competition should dominate interactions among microbial species (9, 11). However, according to the SGH (30), we expect abiotic stress to induce facilitation. Indeed, since MWF is designed to be sterile, it contains biocides, making it a tough and stressful environment for bacteria (35, 37). We next asked whether the observed positive interactions were due to the toxicity of MWF.

### A resource-explicit model predicts that positive interactions occur in toxic environments

To explore the possibility that interactions were due to toxicity, we constructed a mathematical model that describes interspecies interactions through their common exposure to nutrients and toxins in batch culture (Fig. 2B). Our model extends MacArthur’s consumer-resource model (39). For simplicity, we initially considered two species that share and compete for a single limiting nutrient, and are killed by the same toxin, but do not interact otherwise (see Methods). Species deplete the nutrients as they grow, and can invest a proportion of their growth into producing enzymes that degrade the toxin. To match the experiments, we solved the system of equations for each species in mono- and co-culture with a second species and defined (uni-directional) interactions as the difference between the area under the two growth curves. We then used the model to ask how interactions vary as a function of initial nutrient and toxin concentrations.

**Fig. 2.**
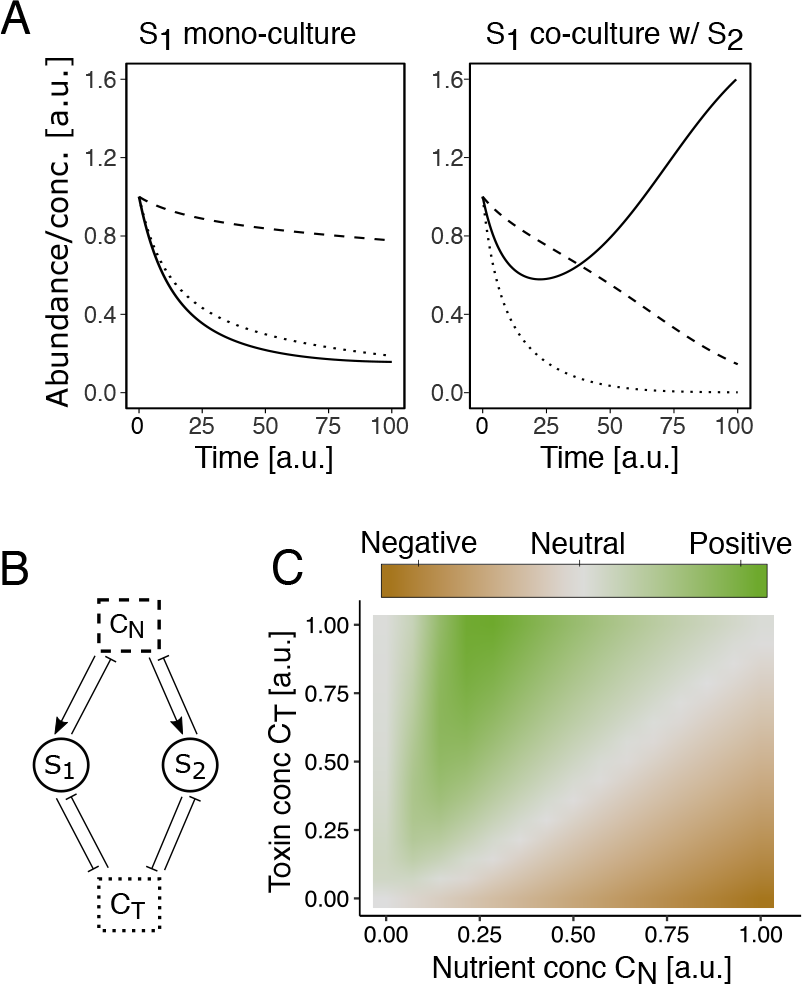
(**A**) Example results of the model (parameters in Table S3), shown as the abundance of species *S*_1_ (solid) and concentrations of nutrients and toxins (dashed and dotted lines). In mono-culture, *S*_1_ goes extinct due to toxins (left), but survives in co-culture with *S*_2_ (right). (**B**) Diagram of our resource-explicit mathematical model where species *S*_1_ and *S*_2_ share a substrate containing nutrients and toxins at concentrations *C*_*N*_ and *C*_*T*_. The species take up the same nutrients, invest a fraction of these into toxin degradation and the rest into population growth. Toxins cause cell death and population decline. (**C**) The response of one species to the presence of another is measured as the difference in AUC between the co- and mono-culture (color, parameters in Table S3) and shown as a function of nutrient and toxin concentrations. At high toxin concentrations and intermediate nutrients, interactions are positive due to the joint degradation of toxins (as in B). As nutrients are increased or toxins decreased, competition for limited resources dominates.

If nutrients are low and toxicity high, species in the model die out regardless of whether they are in mono- or co-culture (grey area on far left of Fig. 2C). As nutrients are increased, the co-cultured species manage to degrade the toxins sufficiently, while bacteria in mono-culture cannot survive (Fig. 2A). In this area of the state-space (green area in Fig. 2C), the presence of the second species has a positive effect on the first (rescuing it from death) despite the underlying competition for nutrients. As nutrients are further increased, however, growth rates increase and toxins can be degraded sooner, such that the presence of a second species becomes unnecessary and even detrimental to the first. The lower the toxin concentration, the faster this competitive effect arises (Fig. 2C). In sum, high toxicity and intermediate nutrients, where species cannot survive alone, is where species in our model benefit from the presence of others. We hypothesized that this regime best describes the four species’ growth in MWF. When the two species have the same model parameters, positive interactions rely on the co-culture being inoculated with twice as many cells as the mono-culture, hence twice the degradation effort. According to our experiments, however, positive interactions still dominate even if the total cell number at the beginning is constant, suggesting that facilitation occurs because different species degrade different toxins (Fig. S5). To better represent this effect, we extended our model in Supplementary Note S2 by introducing a second toxin, and letting each species degrade one of the two. In this extended model, as in the experiments, positive interactions arise even when the total cell number is constant.

### The effect of environmental changes on interspecies interactions matches model predictions

In the model, positive interactions dominate at high toxicity, given that sufficient nutrients are present. Increasing nutrient concentrations further or reducing toxicity instead increase competition. We assumed that our bacteria in the MWF medium lay at the point in the state space where positive interactions are favored, and modified the environment in three additional experiments to test the predictions of the model.

We first increased the concentration of nutrients in the MWF medium by adding 1% amino acids (see Methods), which is a nutrient source for three out of the four species (Fig. S8). In this supplemented MWF medium (MWF+AA), mono-cultures of *A. tumefaciens* and *C. testosteroni* immediately grew well, while *M. saperdae* and *O. anthropi* still suffered from its toxicity (Fig. S9). According to the model, we expect competition between the two species that could grow. Indeed, the two-way positive interaction between *C. testosteroni* and *A. tumefaciens* switched to negative in one direction (Fig. 3B), indicating that a change in nutrient composition can radically modify bacterial interactions. The two species that still experienced the environment as toxic (*M. saperdae* and *O. anthropi*) became the only two species benefiting from being in pairwise co-cultures.

**Fig. 3.**
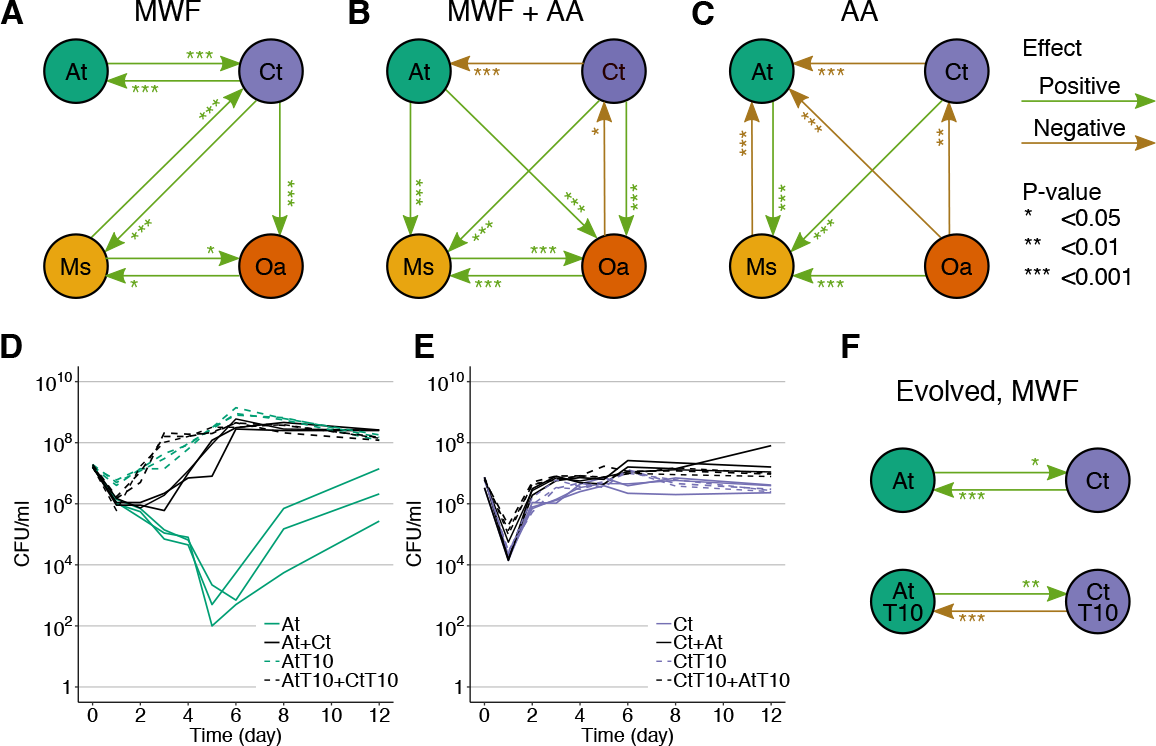
Pairwise interaction networks under different environmental conditions. Positive/negative interactions indicate that the species at the end of an arrow grew significantly better/worse in the presence of the species at the beginning of the arrow in (**A**) MWF, (**B**) MWF+AA and (**C**) AA medium. Statistical significance was calculated based on two experiments in panel (A) (data in Fig. 1 and S1, and one experiment for panels (B) (Fig. S9) and (C) (Fig. S8). All p-values are listed in Table S2. (**D**) Growth curves of ancestral *A. tumefaciens* (At) and (**E**) *C. testosteroni* (Ct) versus the same strains after they had evolved in mono-culture for 10 weeks (AtT10, CtT10). (**F**) Interactions between ancestral and evolved At and Ct strains based on growth curves in panels (D) and (E). The interactions between At and Ct in panels A and F have different p-values because they come from different experimental repeats.

In the second experiment, we reduced the toxicity of our growth medium by growing the bacteria in 1% amino acids (AA). Ideally, we would have removed some of the toxic compounds in MWF, but MWF is chemically complex and is only sold as a finished product. By removing MWF entirely, the growth medium was no longer toxic, but also lacked some of the nutrients in MWF. Caveats aside, according to the model, we expected this change to increase negative interactions. Indeed, we found all interspecies interactions to be negative, except for *M. saperdae*, whose growth was significantly promoted by all three remaining species. *M. saperdae*’s inability to grow in mono-culture in AA (Fig. S8C) suggests that it relies on cross-feeding from the other three species. While our mathematical model does not explicitly capture cross-feeding interactions and assumes that all species compete for the same nutrient, such positive interactions are common in microbial communities (16).

A final way by which we simulated a reduction in environmental toxicity was to allow the bacteria to individually adapt to MWF. We reasoned that if the species evolved to sustain their own growth in MWF, they would lose their positive effects on one another. To test this hypothesis, we conducted experimental evolution on *A. tumefaciens* and *C. testosteroni* by passaging each species alone in MWF for 10 weeks (see Methods, Fig. S10). We did not do this for *M. saperdae* and *O. anthropi* because they could not grow alone in MWF (Fig. 1C, D). After 10 weeks, *A. tumefaciens* grew significantly better in MWF, suggesting that it evolved to become more tolerant to its toxicity (Fig. 3D). In the model, this represents a reduction in toxicity. By again comparing mono- and co-cultures, we found that the positive effect of *C. testosteroni* on *A. tumefaciens* in the ancestral strains switched to competitive in the evolved strains, as predicted by the model (Fig. 3D-F).

Taken together, these results show that positive interactions in our system were most common at high levels of abiotic stress and intermediate nutrient concentrations where most species could not grow, while making the environment more habitable promoted competition. This observation is in line with the SGH. We next took advantage of our system to ask how interactions change with increasing community size.

### Interactions between more than two species depend on environmental toxicity

Our model predicts how the sign of interactions changes with respect to increasing species numbers: in a benign environment with low toxicity, a focal species should grow worse with increasing species number (competition, Fig. 4A). When the number of species is increased in a stressful environment, the increased degradation effort first leads to facilitation. But when enough (functionally equivalent) species are present to alleviate the stress, competition should begin to dominate once again, leading to a hump-shaped curve (Fig. 4A, medium toxicity). This competition arises in the model because all species consume the same nutrient, and would be predicted for communities composed of species whose niches overlap. The community size at which species benefit most from the presence of others (the optimal number of species) depends on the environment as shown in Fig. 4B.

**Fig. 4.**
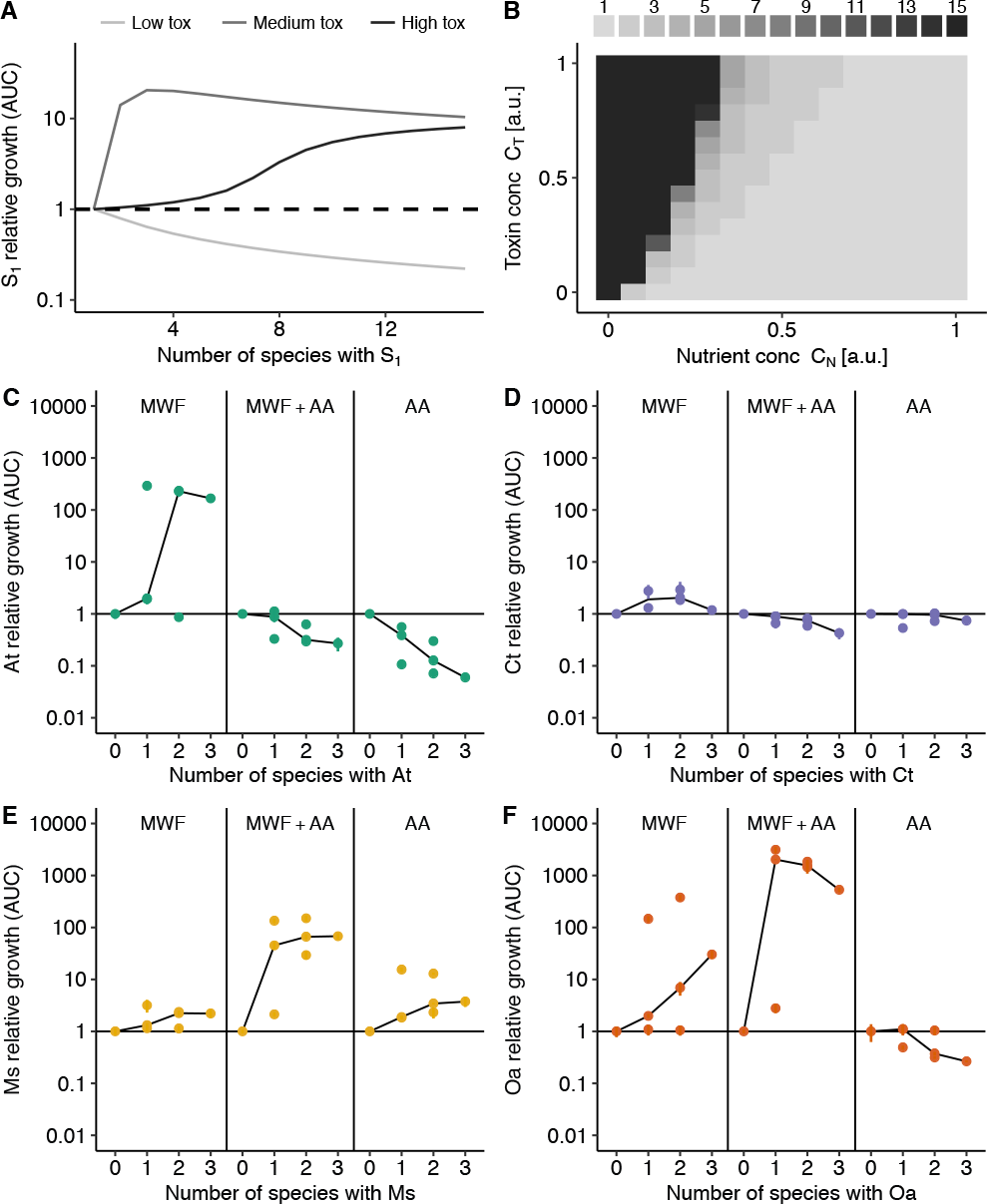
(**A**) Our model predicts that for a focal strain, an increasing community size eventually becomes detrimental. The number at which such competition starts depends on environmental toxicity. (**B**) The optimal number of species with respect to the AUC of a focal strain (peak in panel A) varies with nutrient and toxin concentrations. (**C**–**F**) Each species’ growth expressed in fold-change in its AUC divided by its mean mono-culture AUC in the three different media. Each point shows the mean of a culture treatment composed of 1 to 4 species, and vertical lines show standard deviations. Black lines connect the median points. In environments where a species could not grow alone, the curves are hump-shaped, while in more benign environments, species grow less well in the presence of others.

To test these predictions, we pooled our data from the mono- and pairwise co-cultures (Fig. 1) with experiments where we grew our species in groups of three and four in all three media and calculated the AUC (Fig. S11–S13). In MWF, all species grew better as community size increased (AUC up to 422-fold higher than mono-culture, Fig. 4C-F, left panels). However, this benefit leveled off eventually, resulting in hump-shaped or saturating curves. In MWF+AA, only *M. saperdae* and *O. anthropi*, the two species that couldn’t grow in this medium alone, showed a hump-shaped curve, while *A. tumefaciens* and *C. testosteroni* grew worse with increasing species number. Interestingly, the benefit to *M. saperdae* and *O. anthropi* was considerably higher in MWF+AA than in MWF. This is may be because *A. tumefaciens* and *C. testosteroni* detoxify the environment even further or faster if they can grow well (32, 40, 41). Finally, in AA, increasing competition was observed for all except *M. saperdae*, which was unable to grow alone (Fig. S8C).

In sum, positive interactions occurred in environments that were highly stressful for a species when alone. As this stress was reduced either through the presence of other detoxifying species, or due to increased nutrients or decreased toxicity, competitive interactions between them became salient.

### Degradation efficiency only correlates with species number in toxic environments

Finally, we asked how community size affects its degradation ability and whether that depends on the interactions between its members. In MWF, where interactions were positive (Fig. 3A, 4C-F), increasing species led to better degradation, but did not improve significantly once three species were present (Fig. 5A, F-test comparing the 3-species community with the highest average AAC to the AAC of the 4-species community, *P* = 0.96). Instead, in MWF+AA, where *A. tumefaciens* and *C. testosteroni* experienced competition when other species were added (Fig. 4C, D), degradation efficiency already reached its maximum with a single species, and did not significantly improve in a larger community (*P* = 0.74 for F-test comparing AACs of the communities with the highest average AAC for each community size). Regardless of whether we added AA to the medium, however, a similar final amount of undegraded carbon remained in the four-species communities (Fig. S15). Interestingly, the total population size already saturated at two species in MWF (Fig. S16), suggesting that the benefit in degradation efficiency of a third species is not only due to a larger population size.

**Fig. 5.**
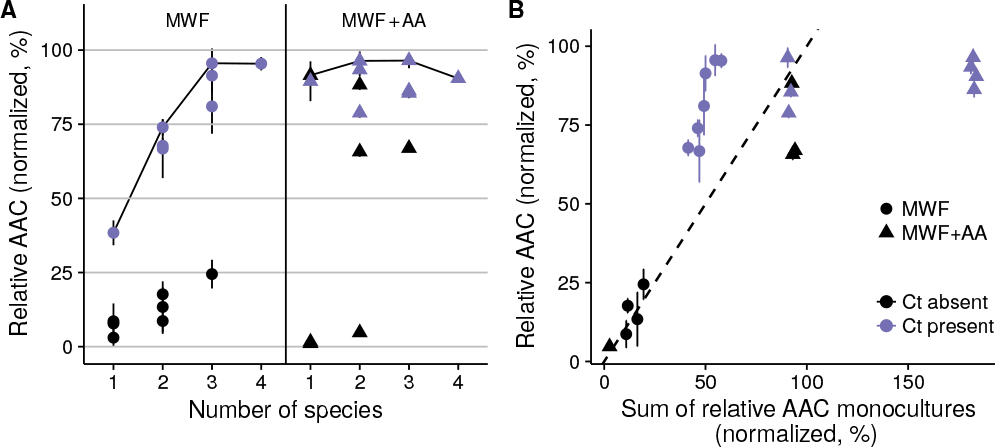
Degradation efficiency as a function of species number. (**A**) Area Above the Curve (AAC) of COD (see Methods), normalized such that all values are between 0 and 100%. Each point shows the mean of a culture treatment composed of 1 to 4 species, and vertical lines show standard deviations. Blue (or black) data-points show cultures where *C. testosteroni* was present (or absent). Cultures growing on MWF (left) only reach their maximum degradation potential once three species are present (see black line connecting the maximum mean values). In MWF+AA (right), even single species can degrade as efficiently as the best cultures. In a more benign environment, there is less need for a diverse community. (**B**) Prediction of an additive model of the sum of degradation efficiencies of individual species is plotted against degradation efficiency of the co-cultures in both growth media. Data-points correspond to co-culture means, and vertical lines show standard deviations (same as data for >1 species in panel (A)). In MWF, co-cultures are more efficient than the sum of the corresponding mono-cultures (most points above dashed line), while in MWF+AA, they are equally or less efficient (most points below the dashed line). The presence of *C. testosteroni* explains much of the AAC in both panels.

The contrast between the two media becomes even clearer if we apply an additive null model to degradation efficiency (i.e., degradation of each species is independent of the other): does the sum of mono-culture degradation efficiencies predict that of the corresponding co-culture? In line with the observed interactions, co-cultures growing in MWF degraded better than the sum of their mono-cultures, while if amino acids were added, the benefit of additional species became minimal (Fig. 5B). A similar analysis on 72 strains (11) found that only few species pairs showed greater productivity in co-culture relative to the prediction of an additive model. Using the same model here, we show that co-culture productivity (i.e. degradation efficiency) changes from being greater to smaller than the null model prediction by simply changing nutrient concentrations.

## Discussion

Quantifying interactions in natural microbial communities remains challenging. By disentangling interactions in a small community, we hope to develop a fundamental understanding that can later be extended to larger ones. What we found in our model system is that facilitative interactions between species occurred in a toxic environment, where only few community members could survive. By presumably improving the environment for their own survival, these species may have accidentally allowed each other to thrive. Once conditions were sufficiently benign, however, competition dominated. These data are in line with the SGH and provide an intuitive explanation for it. Our model (Fig. 4B) then predicts that more diverse communities should be found in toxic environments (40, 42), where similarly, species invasion might be more likely (14, 30).

One important caveat is that we do not know the molecular mechanisms behind the interactions in our system or the process of MWF degradation. These may be important for predicting its behavior. For example, whether degradation occurs through the passive uptake of toxins or through extracellular enzyme secretion will alter predictions on evolutionary stability. It is also unclear why there was a significant decrease in *C. testosteroni*’s population before exponential growth (Fig. 1B). Our model assumes that cells start to grow when enough toxins have been degraded, but it may instead have been because of slow changes in gene expression patterns, or phenotypic heterogeneity in the population (43, 44). Finally, we cannot be sure that facilitation occurs through toxin degradation. However, the positive effect of *C. testosteroni* on many other species (Fig. 3A, S2) suggests that facilitation occurs through the removal of a toxic compound rather than the secretion of a metabolite that so many different species would benefit from.

Nevertheless, our data help address our original question: what makes species in microbial communities help or harm each other? In all the environments where our species could grow, they competed with one another, suggesting that competition is the underlying dynamic between them. Positive effects were instead only observed when species were unable to survive or grow alone. Whether to describe these interactions as cooperative is debatable. A conservative, evolutionary definition of cooperation requires that the relevant phenotype is selected for because of its positive effect on other species (8, 9, 11, 13). Since we have no information on the evolutionary history of the observed behavior, we prefer to refer to it as facilitation (14, 45, 46) and assume that the interactions are an accidental side-effect of each species detoxifying the MWF for its own survival.

The idea that interactions between species can change radically depending on the environment is not new (23–29). Yet current methods to measure and predict interactions in microbial communities lack explicit formulations of context-dependency (15, 47–50). By quantitatively assessing how interactions change with environmental parameters such as substrate concentrations or temperature, we can aim to additionally manipulate community dynamics by carefully engineering the environment (51).

Another major debate in current ecology is whether higher order interactions (HOIs) play an important role in community dynamics (52–57). If HOIs are present, overall community dynamics cannot be predicted based on measurements of interactions between subsets of species within it because the addition of species to these subsets modifies previously measured interactions (58). While we do not explicitly search for HOIs here, we provide a logical argument as to why they may be unavoidable: since each new species added to a community is likely to modify the concentrations of nutrients and toxins, and we know that these concentrations can alter interactions between species pairs (Fig. 2C), then new species can surely modify existing interactions as described by phenomenological models (27, 57). This does not necessarily mean that community dynamics are unpredictable, however, but simply that all components of a system – including substrate concentrations and uptake rates – need to be considered to make accurate predictions. This is difficult in practice, but our argument highlights the need for more mechanistic, resource-explicit models in ecology (28, 59–62).

There is an increasing interest in engineering synthetic microbial communities for practical applications (4, 6, 7, 33, 59, 63, 64). It has been commonly observed that community function saturates with increasing species diversity (64–66). Here we have shown that the rate at which our function of interest (MWF degradation efficiency) saturated depended on environmental toxicity (Fig. 5). This suggests that a harsh environment might require a larger community whose members can facilitate each other’s growth to achieve the desired task. In contrast, making the environment too permissive can reduce the potential benefits of increasing community size due to competition or even competitive exclusion arising between its members. Designing stable consortia in environments where many species are able to grow may therefore be difficult. In other applications, such as antibiotic treatment, where the goal is to eliminate a pathogenic species, it may be that antibiotic toxicity inadvertently leads to facilitation between the surviving organisms. Indeed, we know that antibiotic-resistant bacteria can protect neighboring cells from antibiotics (67–70).

One of the major challenges in current microbial ecology lies in quantifying interactions between species, determining how they are mediated and how they affect community function (2). Using an accessible model system where individual populations and overall community function can be quantified over time has allowed us to address some of these questions. Ecosystems such as this one that use natural bacterial isolates (15, 48, 70–73) are powerful tools that will help disentangle the complexity of natural microbial communities.

## Materials and Methods

### Bacterial species and growth media

This study included four bacterial species: *Agrobacterium tumefaciens*, *Comamonas testosteroni*, *Microbacterium saperdae* and *Ochrobactrum anthropi*. *A. tumefaciens* was modified with a Tn7 transposon containing a GFP marker, and *O. anthropi* with a Tn5 transposon containing an mCherry marker to allow us to distinguish colonies of all four species (see below). The four bacterial species were isolated from waste MWF in a previous study, based on their ability to degrade different MWF substrates (34, 74). It should be noted the waste MWF is less toxic than the fresh MWF that we are preparing here. The identities of the four species were confirmed through 16S gene sequencing. Six additional species isolated from MWF and kindly donated by Peter Küenzi from Blaser Swisslube AG, were also used for supplementary experiments: *Aeromonas caviae*, *Delftia acidovorans*, *Empedobacter falsenii*, *Klebsiella pneumoniae*, *Shewanella putrefaciens*, *Vagococcus fluvialis*. The species were identified at Blaser Swisslube AG by MALDI-TOF, and confirmed by PCR amplification and 16S gene sequencing.

The Metal-Working Fluid (MWF) used in this study (Castrol Hysol™ XF, acquired in 2016) was chosen because of the ability of the four-species co-culture to grow in it and degrade it. The MWF medium was prepared at a concentration of 0.5% (v/v), diluted in water with the addition of selected salts and metal traces to support bacterial growth (Table S1). In addition to (i) the MWF medium, we also conducted growth experiments in (ii) the MWF medium supplemented with 1% Casamino Acids (Difco, UK) (MWF+AA) and (iii) the same selected salts and metal traces supplemented with 1% Casamino Acids only (AA). This third medium was identical to the second, except for the lack of MWF. All medium compositions are listed in Table S1.

### Experimental setup

Before each experiment, each of the four species was independently grown in tryptic soy broth (TSB) overnight starting from a single colony (28°C, 200 rpm) in Erlenmeyer flasks (50 ml) containing 10ml of TSB. The next day, the optical density (OD_600_) of the overnight cultures was measured using a spectrophotometer (Ultrospec 10, Amersham Biosciences), and each species was then inoculated at a standardized OD_600_ of 0.05 into an Erlenmeyer flask (100 ml) containing 20ml of TSB and grown for 3 hours (28°C, 200 rpm) to obtain bacteria in exponential phase with a final concentration of approximately 10^6^-10^7^ CFU/ml at the beginning of each experiment. These starting population sizes were quantified through plating on agar (see below).

For monocultures, 200µl of this final TSB culture were harvested for each species and spun down at 10,000 rcf for 5 minutes. For co-cultures, 200µl of the TSB cultures of each species were first mixed together (e.g. for 2 species, the total was 400µl), then spun down. Experiments were also conducted where the total was fixed to 200µl (Fig. S5). The supernatant was discarded and the pellet resuspended in 30ml of growth medium (e.g. MWF medium) in 100ml glass tubes. In most experiments, 15 treatments (mono-cultures, all pairwise, triplet and quadruplet co-cultures) were conducted simultaneously in triplicate to give 45 experimental cultures in addition to a sterile control. All tubes were incubated at 28°C and shaken at 200 rpm for a total of 12 days.

### Quantifying population size

To quantify the population size of each species over time, 200µl were collected on days 1-6, 8, and 12, from each culture tube, serially diluted and plated onto lysogeny broth (LB) agar or trypticase soy agar (TSA) (Difco, UK) plates and incubated at 28°C to count colony-forming units (CFUs). *C. testosteroni* colonies were visible after 24 hours on TSA, while *A. tumefaciens*, *M. saperdae* and *O. anthropi* were visible after 48 hours on LB agar. To distinguish the latter three species when growing in co-culture, in addition to LB agar, cells were also plated onto LB agar plates containing either: (i) 14.25µg/ml of sulfamethoxazole and 0.75µg/ml of trimethoprim to count only *A. tumefaciens* CFUs; (ii) 2µg/ml of imipenem to count only *M. saperdae* CFUs; or (iii) 10µg/ml of colistin to count only *O. anthropi* CFUs. The fluorescent markers further helped to verify our counts on LB agar.

### Quantifying interspecies interactions

To infer interactions between species, we calculated the area under the growth curve (AUC) of each species in mono-culture and in its pairwise co-culture with each of the other 3 species. We repeated the experiment in the MWF medium on two independent occasions, each in triplicate. We used a blocked ANOVA with “experiment” as a random effect to test for significant differences. If the AUC was significantly greater or smaller in a pairwise co-culture (P<0.05), we deemed the interaction to be positive or negative, respectively. Calculated P-values are shown in Table S2. For the other two media (MWF+AA and AA) and the evolved strains, the pairwise co-culture experiments were performed once only, so F-tests were used to calculate which interactions were significant.

### Quantifying degradation efficiency (Chemical Oxygen Demand)

Chemical oxygen demand (COD) was used as a proxy for the total carbon in the MWF. A significant reduction in COD relative to the sterile control was considered as degradation and the Area Above the Curve (AAC, the integral between the control and the biotic curve) represents degradation efficiency. Briefly, 1ml of MWF emulsion was harvested at the beginning of the experiment, and on days 1-6, 8, and 12, centrifuged (16,000 rcf for 15 minutes) to remove suspended cells (we found that cellular material increases the COD, Fig. S7). Centrifugation separated the MWF into two liquid phases. The top phase was carefully pipetted and discarded, while 200 µl of the second phase was added to NANOCOLOR COD tube tests, detection range 1-15 g/l by Macherey-Nagel (ref: 985 038), heated at 160°C for 30 mins, cooled to room temperature, and the color change quantified on a LASA 7 100 colorimeter (Hach Lange, UK).

### Adapting bacteria to MWF medium

*A. tumefaciens* and *C. testosteroni* were grown in MWF medium as described above for 7 days (28°C, 200 rpm) in five replicate monocultures. After 7 days, 30 ml of fresh MWF medium was prepared and 300 µl of the week-old culture transferred into it. This was repeated every week for a total of 10 weeks. At the beginning and at the end of every week, population sizes were quantified using CFUs as described above. After 3 weeks, three replicate populations of *A. tumefaciens* had gone extinct (Fig. S10). After 10 weeks, one colony was isolated from the first replicate of the evolved populations of *A. tumefaciens* and *C. testosteroni*, and the interactions between them quantified.

### Resource-explicit mathematical model

We consider our community to consist of *n* distinct species, where the change in abundance *S*_*i*_ of species *i* is determined by a growth function *ρ_i_* and mortality *µ_i_* which depend on the concentrations *C*_*N*_ and *C*_*T*_ of the nutrient and toxin as shown in Fig. 2B. Nutrient concentrations decrease as a function of the species’ growth via the biomass yield *Y*_*i*_, while toxin concentrations decrease according to the species’ production rate *δ_i_* of enzymes that degrade the toxin as well as a passive uptake rate *κ_i_*. A fraction *f*_*i*_ of the collected nutrients are invested into active degradation and the rest into growth. This results in the following set of differential equations:

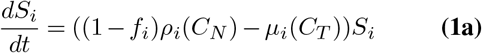

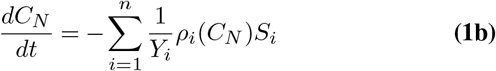

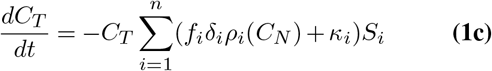

We assume that the growth and death rates saturate with increasing nutrient or toxin concentrations as:

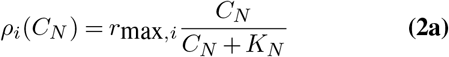

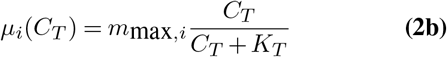

for the nutrient and toxin with half-saturation concentrations *K*_*N*_, *K*_*T*_ and maximum growth or death rate *r*_max_, *m*_max_. We implemented the model in Python v3.6 using the SciPy library v1.0 and solved with standard ODE solvers for a set of parameters and initial conditions as listed in Table S3. Fig. S17 shows how changes in these parameters and initial conditions affect the outcome of the model. To generate the heat plot in Fig. 2C, we calculated the difference in the AUC of the simulated time-series, between a simulation with initial abundance *S*_1_ = *S*_2_ = 1 and another with initial abundance *S*_1_ = 1 and *S*_2_ = 0.

## Supporting information

Supplementary figures and model extension

## ACKNOWLEDGEMENTS

We thank Jake Alexander, Kevin Foster, Laurent Keller and all members of the Mitri lab for useful discussions and feedback on the manuscript. We thank Christopher van der Gast and Ian Thompson for providing the four strains used in the study, and for their advice in establishing the protocols. We thank Peter Küenzi and for providing the additional six MWF isolates, and to Sabrina Rivera for conducting growth experiments with them. We thank Guillermo Osuna for help developing the COD measurement protocol, Marc Garcia-Garcerà and Bastien Vallat for performing colony PCRs and 16S sequencing of the six additional strains, and Avery Becker for producing the dead cells. PP conducted experiments, BV developed the mathematical model, and SM conceived of and supervised the project. All authors participated in the writing of the manuscript. PP is funded by the University of Lausanne. BV and SM are funded by European Research Council Starting Grant 715097.

