## Supplementary figures and model extension for "Toxicity drives facilitation between four bacterial species"

### Supplementary Note S1: Figures

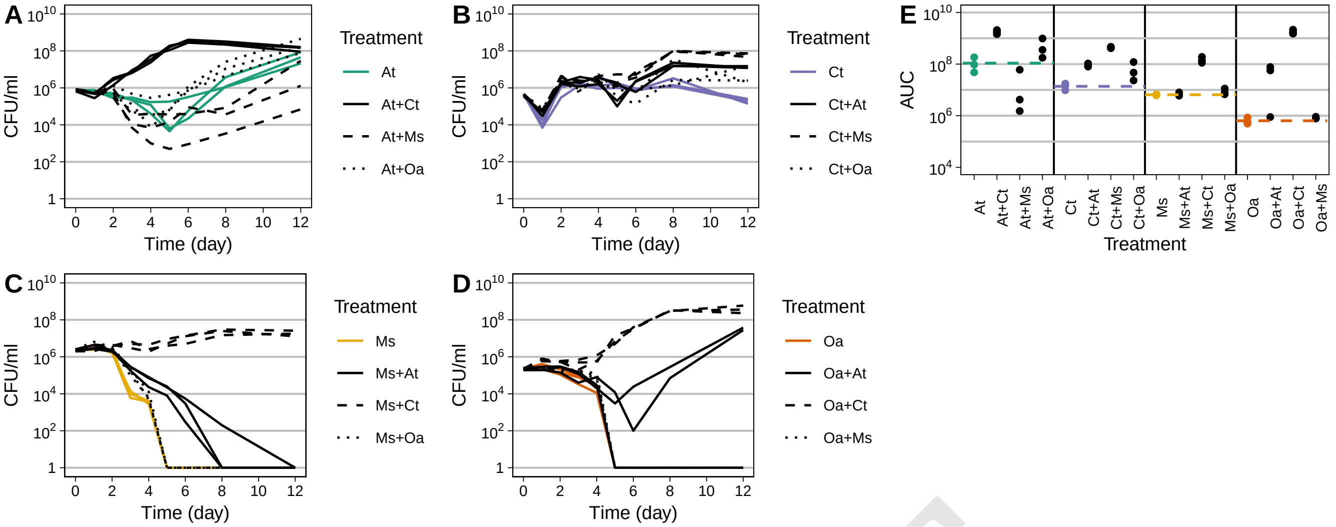


**Fig. S1.** Repetition of the experiment shown in Fig. 1. At this point in time our protocol for measuring the COD was not yet optimized, so we do not show those data. The AUC data are combined with the data in Fig. 1E to perform the statistical test.

DRAFT

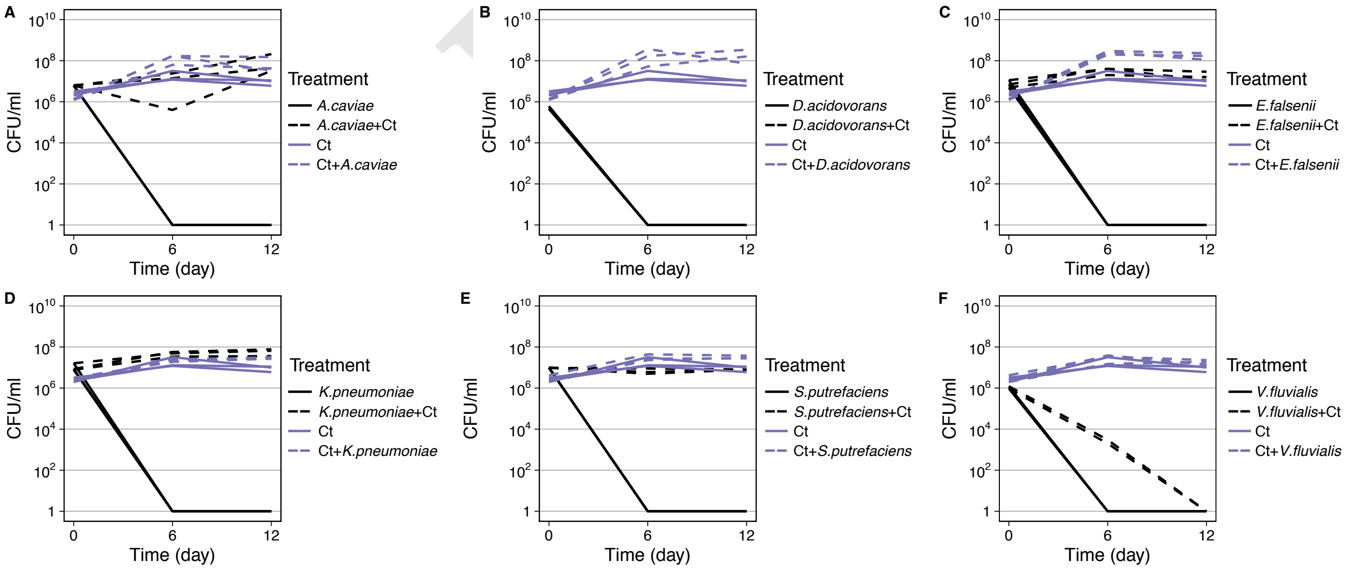


**Fig. S2.** Pairwise co-cultures in triplicate of *C. testosteroni* (Ct) together with six other strains that were similarly isolated from MWF, but had presumably never interacted previously with *C. testosteroni*. In co-cultures, the strain mentioned first in the label is the one plotted (e.g. Ct + *A. caviae* is the growth curve of Ct in the presence of *A. caviae*). All six strains quickly died in MWF if cultured alone, but in the presence of *C. testosteroni*, four out of the six grew better, as shown in Fig. S3. This suggests that the positive interactions at least between *C. testosteroni* and the other three strains shown in Fig. 3 are unlikely to be a product of their co-evolution, but are most likely accidental. Species identities are: (A) *Aeromonas caviae*, (B) *Delftia acidovorans*, (C) *Empedobacter falsenii*, (D) *Klebsiella pneumoniae*, (E) *Shewanella putrefaciens*, (F) *Vagococcus fluvialis*.

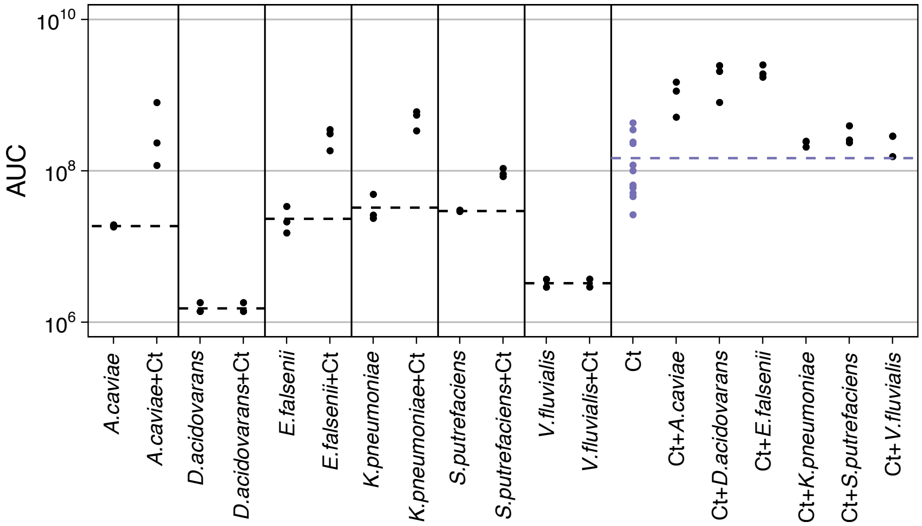


**Fig. S3.** AUC calculated based on the data in Fig. S2. Four out of the six strains had a higher AUC when grown together with *C. testosteroni*, while *C. testosteroni* also improved its growth in some cases. Note that a mono-culture of *C. testosteroni* was not included every time, but a number of repeats of its AUC are pooled together. This makes a statistical comparison for Ct more difficult, but supports our overall conclusion that positive interactions are likely accidental.

DRAFT

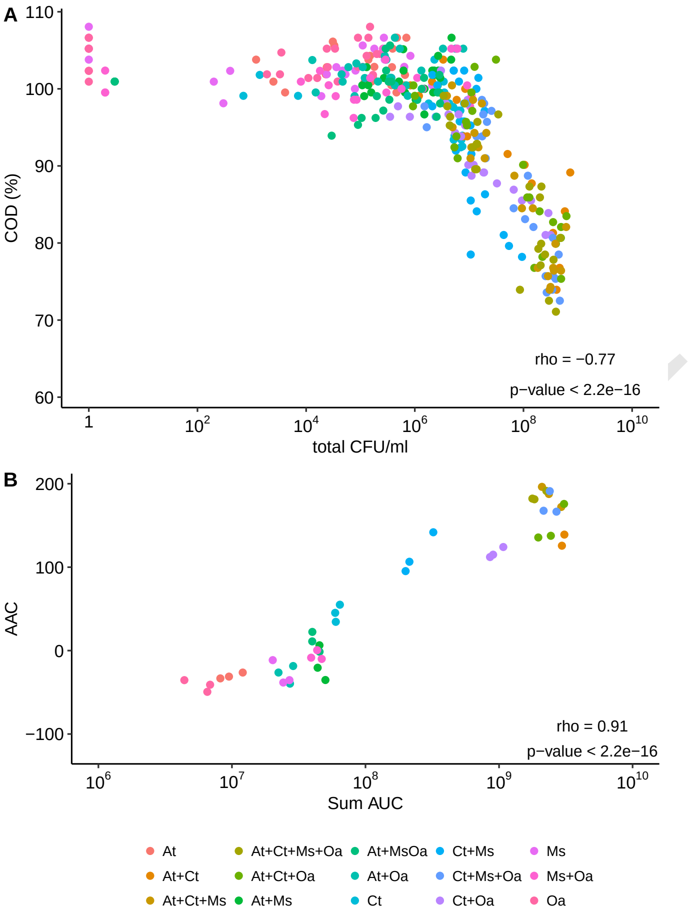


**Fig. S4.** (A) CFU/ml and COD correlate across experimental conditions (Spearman's correlation test). Data from time-point 0 were not included in this analysis. (B) AUC also correlates with AAC (Spearman's correlation test).

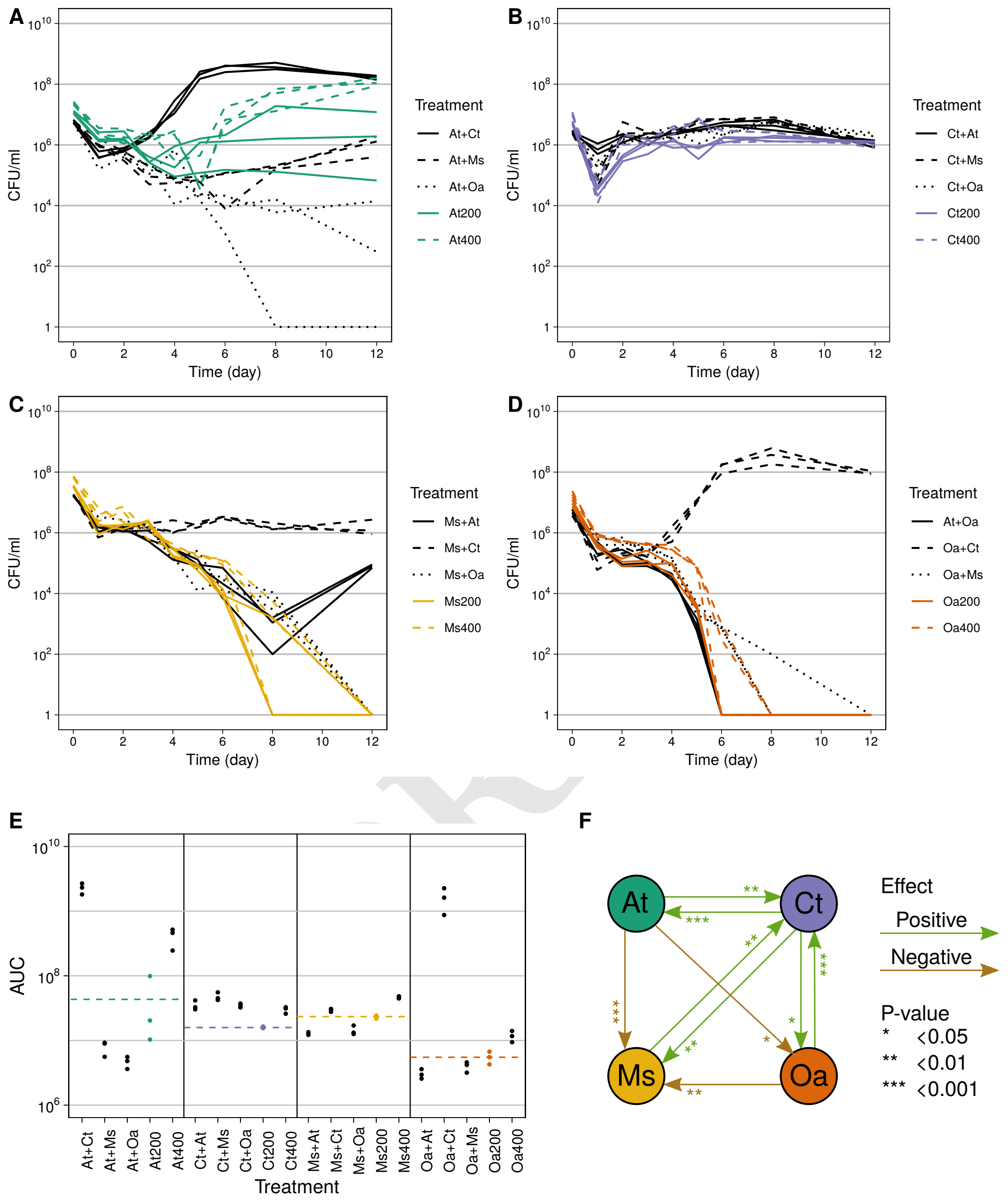


**Fig. S5.** Repeat of the mono-culture and pairwise co-cultures presented in the main text (Fig. 1) but where co-cultures contained the same total inoculum volume as the mono-cultures (e.g. At200 was inoculated with 200$\mu$l of At, while At+Ct was inoculated with 100$\mu$l of At and 100$\mu$l of Ct). We also included a treatment where we doubled the volume of the inoculum for the mono-cultures (e.g. At400). (A)-(D) Growth curves for each species, (E) AUC for each treatment and species, (F) interaction network comparing mono-cultures and co-cultures of the same total inoculum volume (e.g. comparing At200 with At+Ct of 100 each). For all species, doubling the inoculum volume of the mono-culture resulted in AUCs that were significantly larger (all four $P < 0.015$).

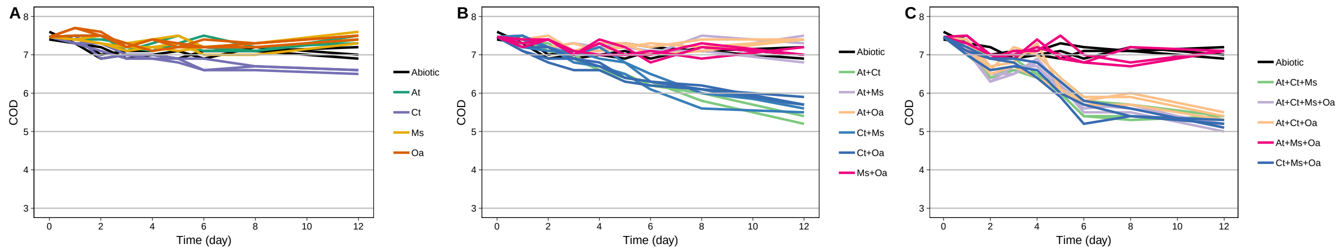


**Fig. S6.** Chemical Oxygen Demand (COD) over time in MWF medium in (A) mono-cultures, (B) pairwise co-cultures and (C) in three- and four-species co-cultures. Each treatment was performed in triplicate. Black lines show an abiotic control treatment that was sterile. Note that these absolute COD values are not directly comparable to those in Fig. S14, since for each experiment the values in the abiotic control differ somewhat.

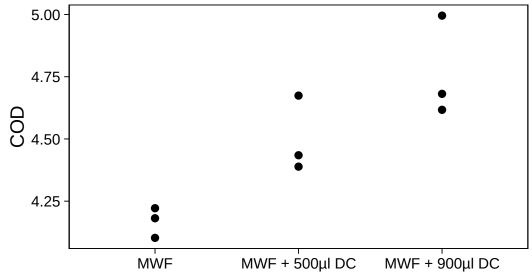


**Fig. S7.** Chemical Oxygen Demand (COD) an abiotic control treatment that was sterile compared to two treatments containing increasing concentrations of dead cells. Cells were killed by autoclaving. This shows that dead cells can increase the COD.

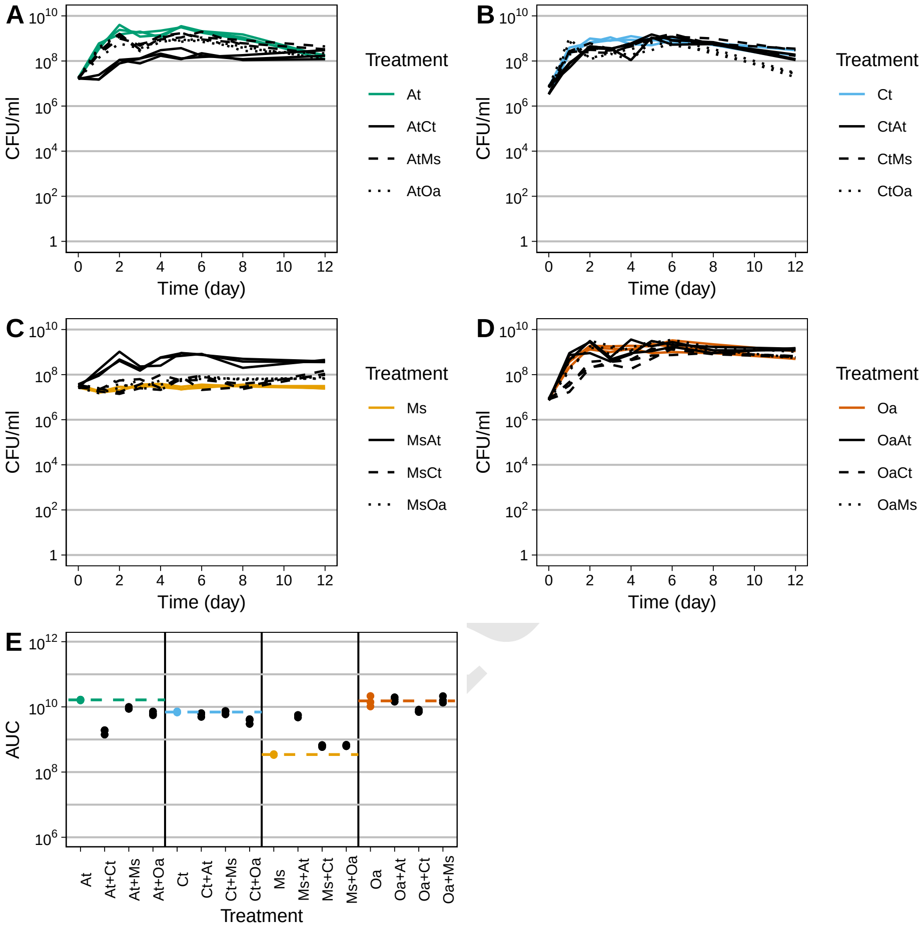


**Fig. S8.** Growth and pairwise interactions in amino acid medium (AA).

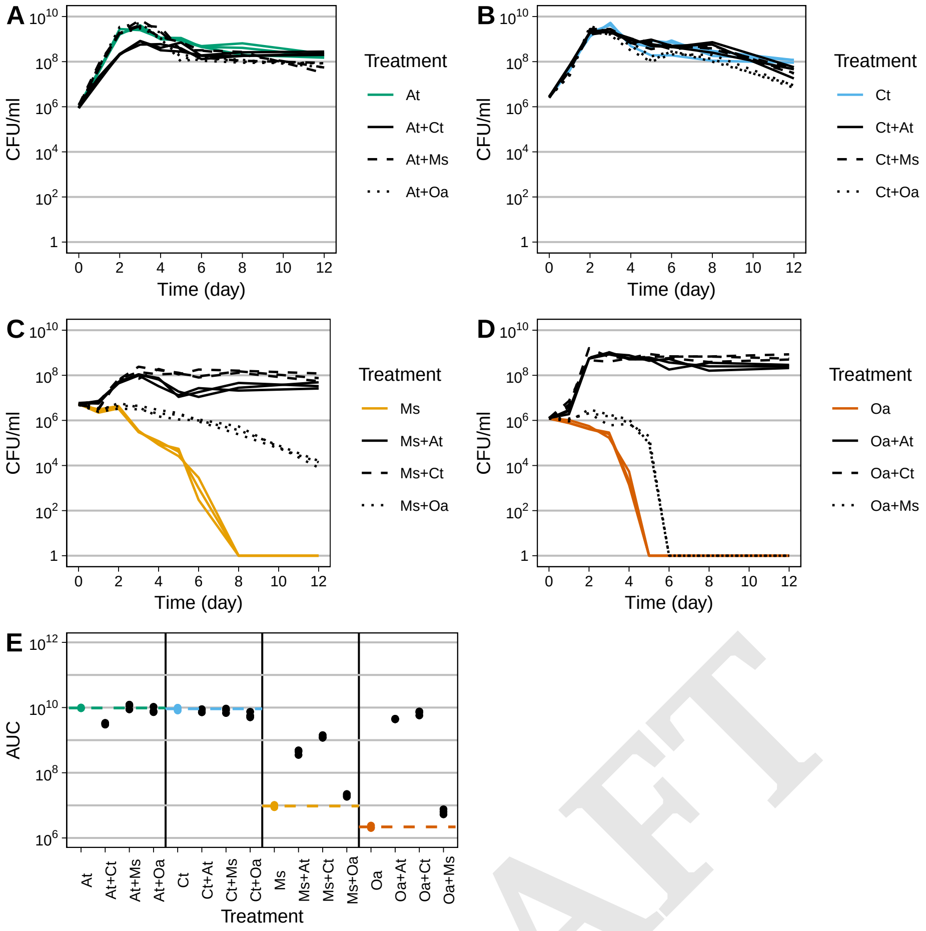


**Fig. S9.** Growth and pairwise interactions in MWF supplemented with amino acids (MWF+AA).

DRAFT

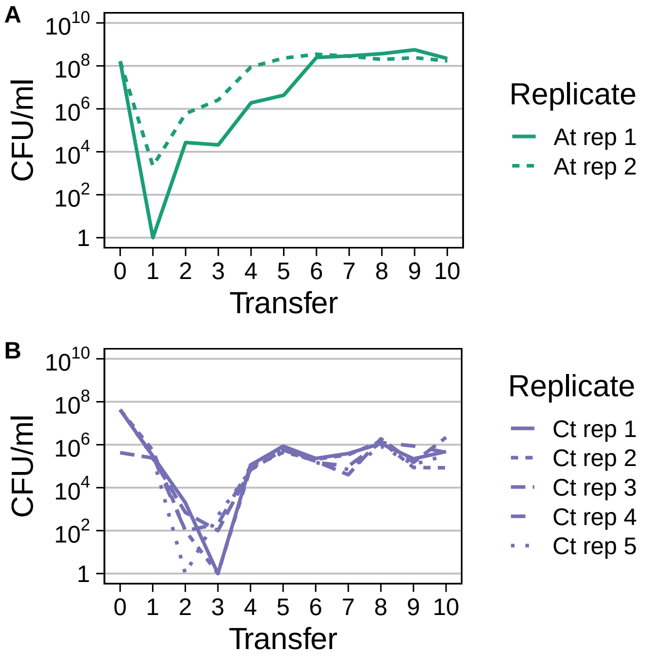


**Fig. S10.** Evolutionary experiment where mono-cultures of (A) *A. tumefaciens* and (B) *C. testosteroni* were transferred into fresh tubes every week. The plot shows CFU/ml at the end of each week prior to a 1:100 dilution. Five replicates were used for each condition, but in three *A. tumefaciens* cultures, cells went extinct after a few weeks. Clones were taken for each species from replicate one (solid line) for the experiments shown in Fig. 3.

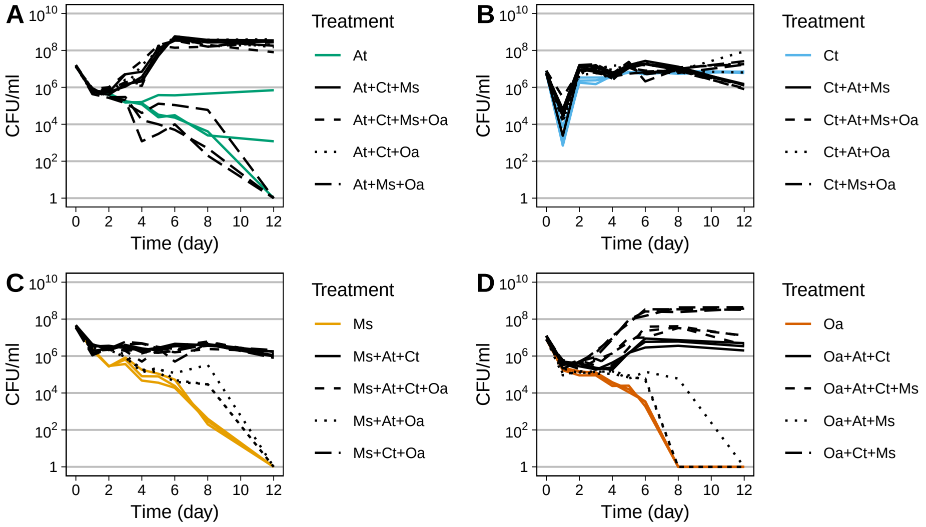


**Fig. S11.** Growth and interactions of three and four-species co-cultures in MWF medium.

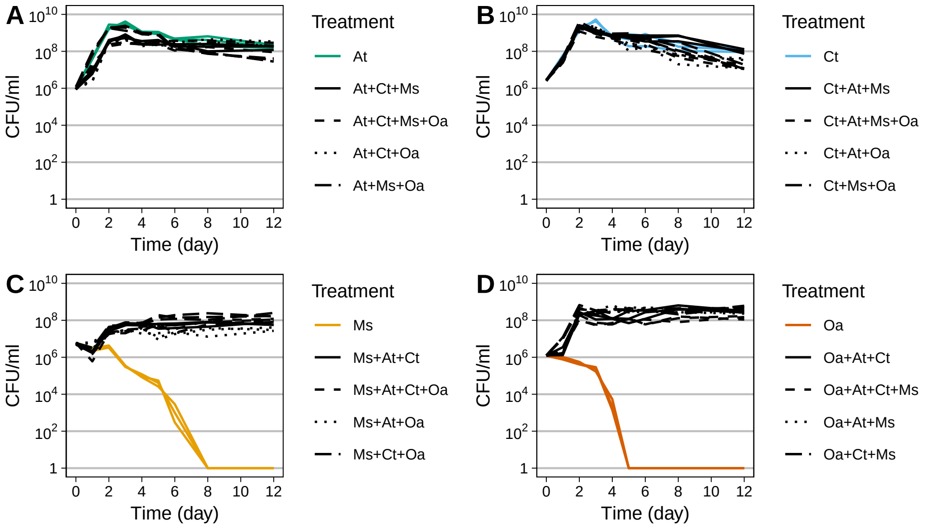


**Fig. S12.** Growth and interactions of three and four-species co-cultures in MWF+AA medium.

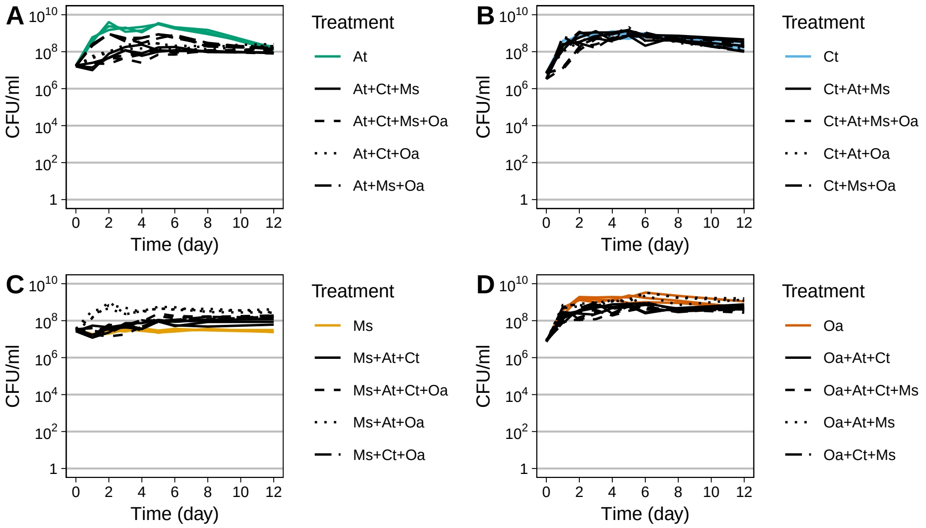


**Fig. S13.** Growth and interactions of three and four-species co-cultures in amino acid medium (AA).

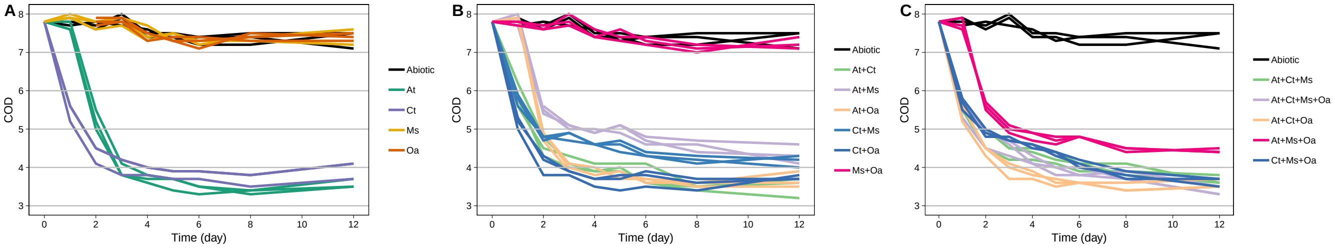


**Fig. S14.** Chemical Oxygen Demand (COD) over time in MWF+AA medium in (A) mono-cultures, (B) pairwise co-cultures and (C) in three- and four-species co-cultures. Each treatment was performed in triplicate. Black lines show an abiotic control treatment that was sterile. Note that these absolute COD values are not directly comparable to those in Fig. S6, since for each experiment the values in the abiotic control differ somewhat.

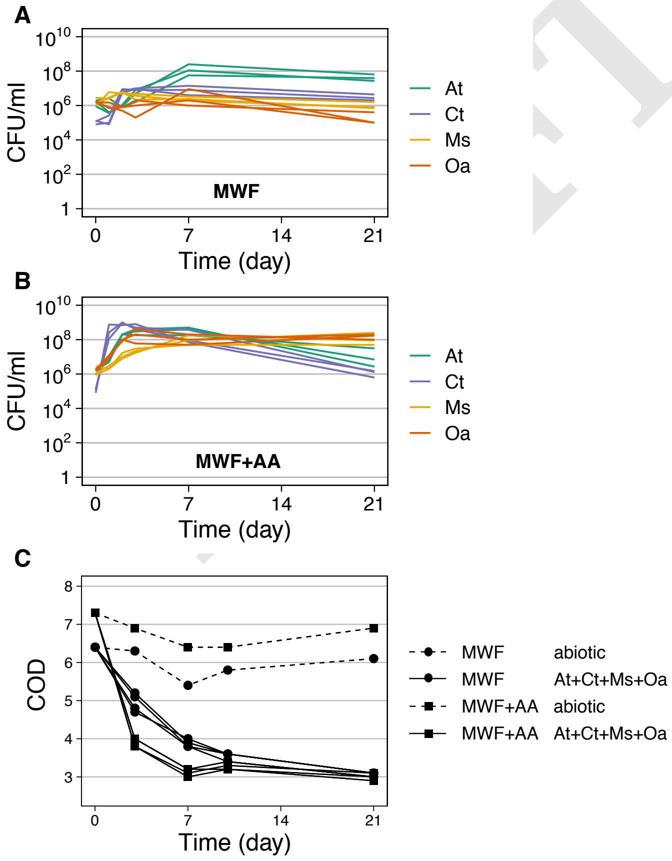


**Fig. S15.** CFU/ml and Chemical Oxygen Demand (COD) over time in an additional experiment where all four species were grown in MWF and in MWF+AA in parallel. (A) Growth in MWF as well as (B) in MWF+AA stabilizes after around 7 days, and matches results shown in Fig. S11 and S12. (C) COD decrease changes little after 12 days corresponding to the length of our experiment. Note that in each experiment, the starting COD of the control is somewhat different, making it difficult to compare COD values between experiments. Here, however, we can directly compare the COD decrease in MWF and in MWF+AA. These data show that AA has a COD of approximately 0.9g/L, and that the four species appear to degrade similar substrates independently of the presence of AA, which can be seen in how the COD converges to a similar value in the two treatments.

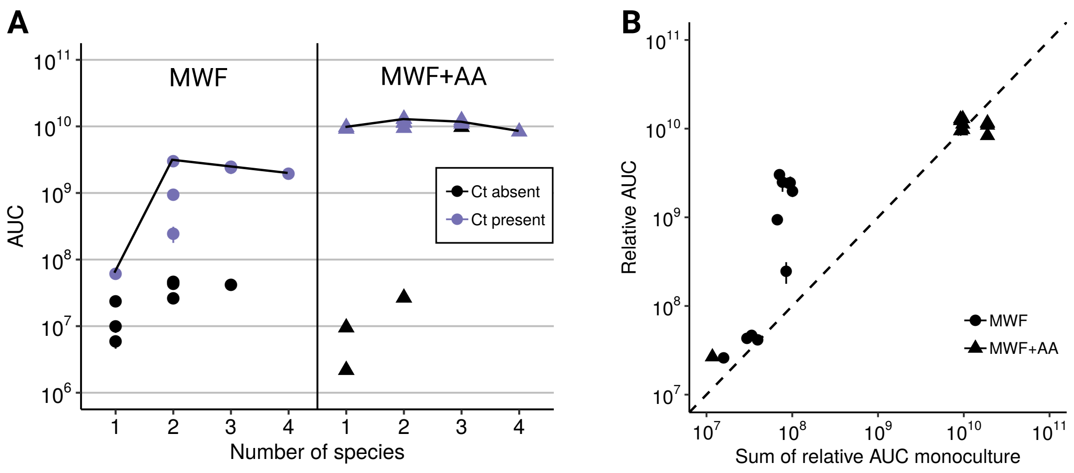


**Fig. S16.** Same as Fig. 5 in the main text but rather than the COD data and the AAC, here we use the total CFU/ml data and the AUCs of each mono- and co-culture to observe at what species number the overall population size saturates. In (A) we ask whether the carrying capacity increases with additional species and find that in MWF it reaches its maximum at 2 species, while in MWF+AA, it is already reached at 1 species. (B) The sum of AUCs in mono-culture do not always predict the co-culture. Again, in MWF, populations are sometimes larger in co-culture than expected by the additive null model, while for MWF+AA, the null model is a good predictor for the total in co-culture. Note that the scale here is logarithmic.

| Medium | Product | Concentration |
|---|---|---|
| MWF | $K_2HPO_4$ | 0.6g |
| | $KH_2PO_4$ | 0.6g |
| | HMB | 20ml |
| | Castrol Hysol™ XF | 500ml |
| | $H_2O$ | to complete 1L |
| MWF+AA | $K_2HPO_4$ | 0.6g |
| | $KH_2PO_4$ | 0.6g |
| | HMB | 20ml |
| | Castrol Hysol™ XF | 500ml |
| | casaminoacids | 0.1g |
| | $H_2O$ | to complete 1L |
| AA | $K_2HPO_4$ | 0.6g |
| | $KH_2PO_4$ | 0.6g |
| | HMB | 20ml |
| | casaminoacids | 0.1g |
| | $H_2O$ | to complete 1L |
| HMB | NTA (nitric triacetic acid) | 10g |
| | $MgSO_4.7H_2O$ | 14.45g |
| | $CaCl_2.2H_2O$ | 3.33g |
| | $(NH_4)_6Mo_7O_{24}.4H_2O$ | 0.00974g |
| | $FeSO_4.7H_2O$ | 0.099g |
| | Metals44 | 50ml |
| | $H_2O$ | to complete 1L |
| Metals44 | $Na_2EDTA.2H_2O$ | 0.387g |
| | $ZnSO_4.7H_2O$ | 1.095g |
| | $FeSO_4.7H_2O$ | 0.914g |
| | $MnSO_4.7H_2O$ | 0.154g |
| | $CuSO_4.H_2O$ | 0.0392g |
| | $Co(NO_3)_2.6H_2O$ | 0.0248g |
| | $Na_2B_4O_7.10H_2O$ | 0.0177g |
| | $H_2O$ | to complete 100ml |

DRAFT

**Table S1.** Chemical composition of media used.

**Table S2.** P-values behind interactions in Fig. 3 are shown in attached Excel file.

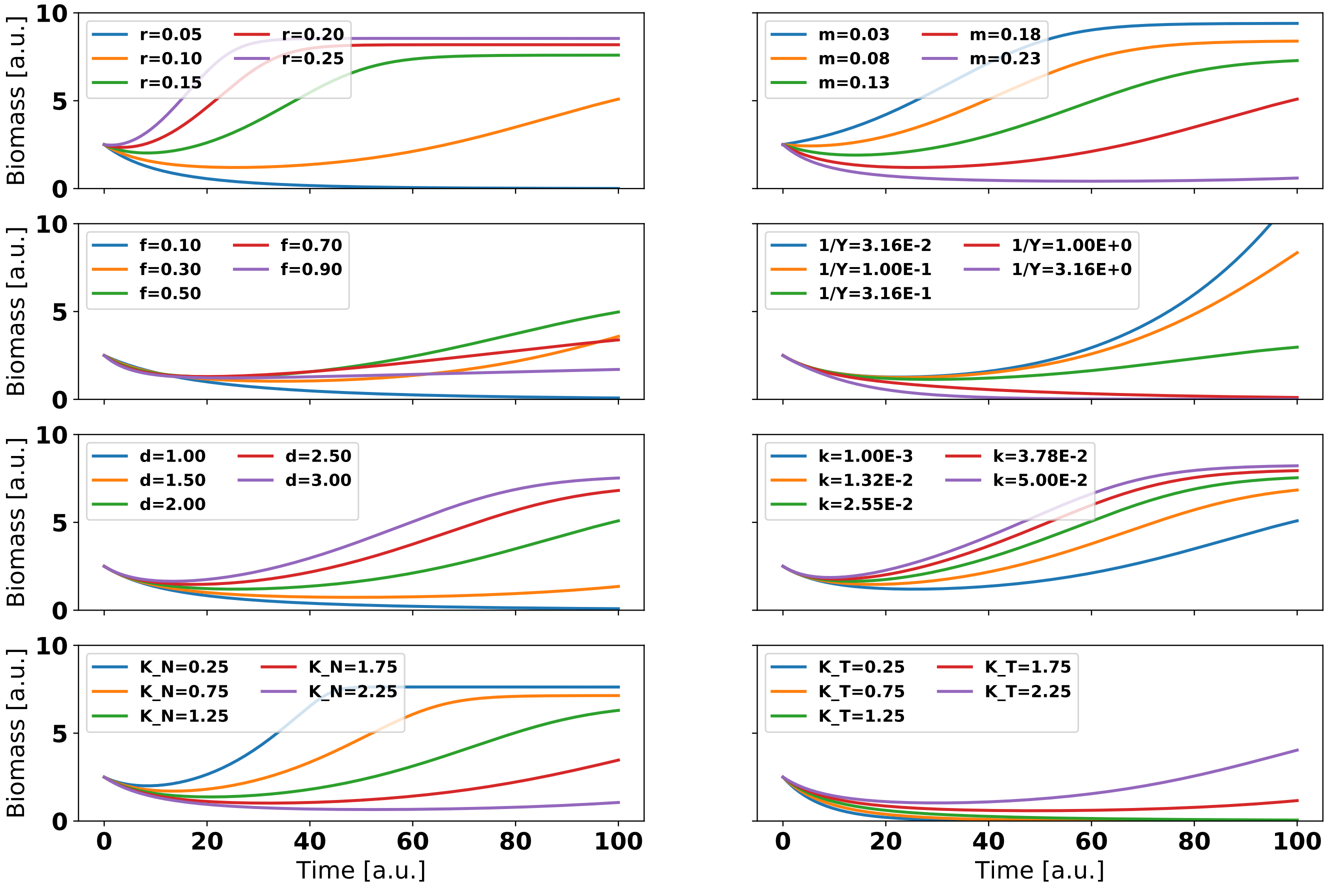


**Fig. S17.** Effect of parameters on simulated monoculture biomass in the mathematical model Eq. (S1a)-Eq. (S1c). Row-wise from top left to bottom right: maximum growth rate $r_{max}$, maximum death rate $m_{max}$, degradation investment $f$, inverse of biomass yield $Y$, degradation rate $\delta$, passive degradation rate $\kappa$, nutrient and toxin saturation constants $K_N$, $K_T$. When one parameter is changed, the rest are kept at the standard value found in Table S3.

| Param | $r_{max}$ | $m_{max}$ | $Y$ | $\delta$ | $\kappa$ | $f$ | $K_N$ | $K_T$ |
|---|---|---|---|---|---|---|---|---|
| Figure 2B | 0.1 | 0.2 | 0.2 | 15.0 | $1.0E-3$ | 0.1 | 1.5 | 1.0 |
| Figure 2C, 4AB | 0.1 | 0.15 | 0.2 | 10.0 | $1.0E-2$ | 0.6 | 1.0 | 1.0 |

**Table S3.** These parameters were used to run the examples in Figures 2B,C and 4A, B in the main text. The parameters were tuned to show representative dynamics of the mathematical model described in Eq. (S1a)–Eq. (S1c).

### Supplementary Note S2: Differential toxin degradation allows for density-independent facilitation

The original formulation in the main text (copied below for ease of reading) is valid for $n$ species affected by a single toxin and sharing a single nutrient. For simplicity, in the main text we assume that the two species are identical in all parameters, while in practice they will probably vary in their mortality rates $m$, toxin degradation rates $\delta$, growth rate $r$ and biomass yield $Y$. Recall that the we use Monod kinetics for the growth function $\rho_i(C_N) = r_i C_N/(C_N + K_N)$, (2A) and toxin-dependent mortality $\mu_i(C_T) = m_i C_T/(C_T + K_T)$, (2B).

$$\frac{dS_i}{dt} = ((1-f_i)\rho_i(C_N) - \mu_i(C_T))S_i \quad \textbf{(S1a)}$$

$$\frac{dC_N}{dt} = -\sum_{i=1}^{n} \frac{1}{Y_i}\rho_i(C_N)S_i \quad \textbf{(S1b)}$$

$$\frac{dC_T}{dt} = -C_T \sum_{i=1}^{n} (f_i\delta_i\rho_i(C_N) + \kappa_i)S_i \quad \textbf{(S1c)}$$

Importantly, we have shown in the main text that both positive and negative interactions may occur in an environment that includes toxins, where the positive interactions relies on increasing the total population size from mono- to co-culture. If instead the total population size is kept constant by inoculating the $n$ species co-cultures with initial abundance $\frac{1}{n}$, only negative interactions will be found, because the same resources will be divided amongst all strains. On the contrary, our experiments revealed positive interactions even when the total population size was held constant (Fig. S5).
To better capture these positive interactions when scaling the inoculum in this manner, we modified the model to include two toxins $T_1$ and $T_2$, which affect both strains equally, as shown in Eq. (S2a)–Eq. (S2e) and Fig. S18A, where $T_1$ is degraded by strain $S_1$ while $S_2$ degrades $T_2$. At intermediate nutrient concentrations, we again observe two-way positive effects of being in co-culture (Fig. S18) since the degrading member of each toxin facilitates the environment for the other by reducing the concentration of one of the two toxins.

$$\frac{dS_1}{dt} = ((1-f_1)\rho_1(C_N) - \mu_1(C_{T_1}) - \mu_2(C_{T_2}))\,S_1 \quad \textbf{(S2a)}$$

$$\frac{dS_2}{dt} = ((1-f_2)\rho_2(C_N) - \mu_1(C_{T_1}) - \mu_2(C_{T_2}))\,S_2 \quad \textbf{(S2b)}$$

$$\frac{dC_N}{dt} = -\left(\frac{1}{Y_1}\rho_1(C_N)S_1 + \frac{1}{Y_2}\rho_2(C_N)S_2\right) \quad \textbf{(S2c)}$$

$$\frac{dC_{T_1}}{dt} = -C_{T_1}(f_1\delta_1\rho_1(C_N)S_1 + \kappa_1 S_1) \quad \textbf{(S2d)}$$

$$\frac{dC_{T_2}}{dt} = -C_{T_2}(f_2\delta_2\rho_2(C_N)S_2 + \kappa_2 S_2) \quad \textbf{(S2e)}$$

DRAFT

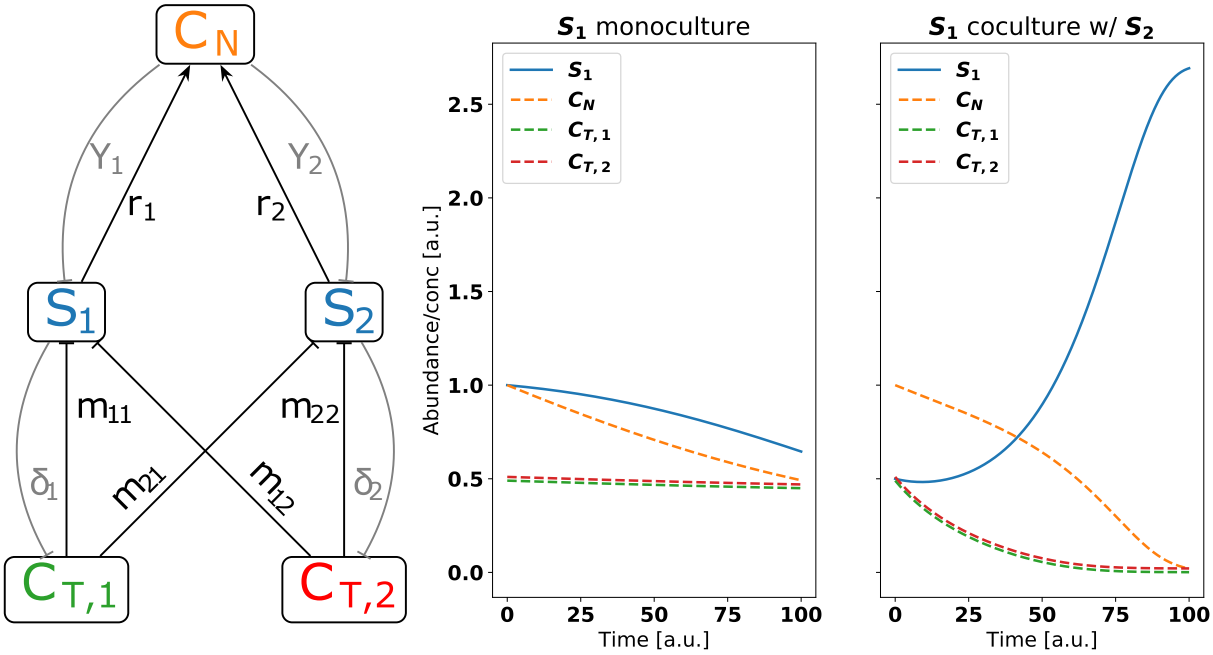


**Fig. S18.** (**A**) We extend the model by introducing two toxins that affect both species but can only be degraded by one species each. (**B**) The simulated biomass shows positive interactions where both species are better off in co-culture even while the total inoculated biomass is the same between mono- and co-culture. In the example run, the effect of nutrients and toxins is symmetric by the choice of equal parameters for $S_1$ and $S_2$, and the two species only differ in their degradation abilities for the two toxins.
